# Efficient colonic colonization by *Campylobacter jejuni* requires the heme receptor ChuA

**DOI:** 10.64898/2026.08.04.742739

**Authors:** Binita Baral, Madison L. Bunch, Emily L. Roberts, Vincent R. Randaisi, Wyatt W. Wittliff, William N. Beavers, Andrew J. Monteith, David K. Meyerholz, Jeremiah G. Johnson

## Abstract

Previous research demonstrated that *Campylobacter jejuni* encodes a heme utilization system that facilitates heme-dependent growth under iron-limiting conditions and that transcription of this system is induced during human infection. Despite these observations, it remained unknown whether the heme transport system is required for colonization and disease in a susceptible host. To address this, we created individual non-polar deletion mutants of each component of the heme transport system and examined their ability to promote heme-dependent growth and iron uptake. From this work, we found that only the heme receptor, ChuA, was required for heme-dependent growth and iron acquisition, which supports the earlier work of another group. Further, we examined whether intestinal colonization, immune activation, and pathology were altered during infection with these mutants. After establishing that elevated heme and *chuABCD* expression occur during *C. jejuni* infection of IL-10^−/−^ mice, we found that a mutant of the heme receptor, ChuA, exhibited significantly reduced colonization of the colon. In addition, we found that neutrophil and circulating monocyte recruitment were significantly reduced in the colon during infection with the ChuA mutant, but that populations of self-maintaining tissue macrophages remained high. Loss of ChuA reduced colonic colonization and was accompanied by diminished innate immune cell recruitment and intestinal pathology. Together, these findings identify ChuA-dependent heme acquisition as a key determinant of efficient colonic colonization and the associated inflammatory disease.

## Introduction

*Campylobacter jejuni* is a leading cause of bacterial gastroenteritis with an estimated 1-2 million infections in the United States and 170 million infections globally [1, 2]. This prevalence is due to the bacterium residing at high numbers in the gastrointestinal (GI) tracts of several animal species, including birds, cattle, and pigs [3, 4]. Infection then occurs through consuming undercooked, contaminated meat products, milk, water, or unwashed produce. Once in the lower gastrointestinal tract, *C. jejuni* invades ileocecal tissues and/or transits paracellularly, which results in an inflammatory gastroenteritis that is characterized by diarrhea, fever, hematochezia, and abdominal pain [5–8]. While infection often resolves without intervention, several post-infectious inflammatory disorders are associated with human campylobacteriosis, including Guillain-Barré Syndrome, inflammatory bowel disease, and reactive arthritis [7–11]. Further, the organism has been designated a serious threat to public health in the U.S. due to its increasing resistance to fluoroquinolones and macrolides [12].

During GI infection, bacterial pathogens often alter their gene expression profiles to successfully colonize a host, including processes involved in counteracting the host’s immune response and acquiring sufficient nutrients to facilitate growth [13–15]. In a previous study that used transcriptomics to identify *C. jejuni* determinants that were altered during human infection, several genes involved in iron acquisition were identified, including some of those in the *chuZABCD* heme utilization system [16]. Similarly, a study using human fecal extracts to identify human-specific transcriptomic changes in *C. jejuni* observed that *chuC* alone was upregulated in the presence of human fecal extracts [17]. Because earlier studies demonstrated *chuZABCD* is under the control of the ferric uptake regulator (Fur) and induced under iron-restricted conditions, the above data suggest that the human gut is limited in free iron and that *C. jejuni* responds by scavenging for iron from alternative sources, such as heme [18].

Acquisition of iron from heme during bacterial infection is well-established for both Gram-negative and Gram-positive pathogens. In Gram-negative pathogens, heme utilization occurs through binding a receptor in the outer membrane with initial transport being driven by the TonB/ExbB/ExbD motor complex [19–22]. Once in the periplasm, heme is often bound by a periplasmic chaperone and guided to a cognate inner membrane ATP-binding cassette (ABC) transport complex, which facilitates translocation into the cytoplasm [19, 20]. In the cytoplasm, a heme oxygenase hydrolyzes the porphyrin ring to release the bound iron so it may be used by the bacterium. *C. jejuni* encodes most of these components in a cluster that contains the heme oxygenase gene, *chuZ*, which is divergently transcribed from an operon encoding the transport determinants: ChuA, the heme receptor, and ChuBCD, the ABC transporter (Fig. 1A) [16]. In previous studies, the heme receptor, the Ton motor complex, and the heme oxygenase were shown to facilitate heme-dependent growth of *C. jejuni* under iron-restricted conditions while the ABC transporter was shown to be dispensable (Fig. 1B) [18, 23].

**Figure 1.**
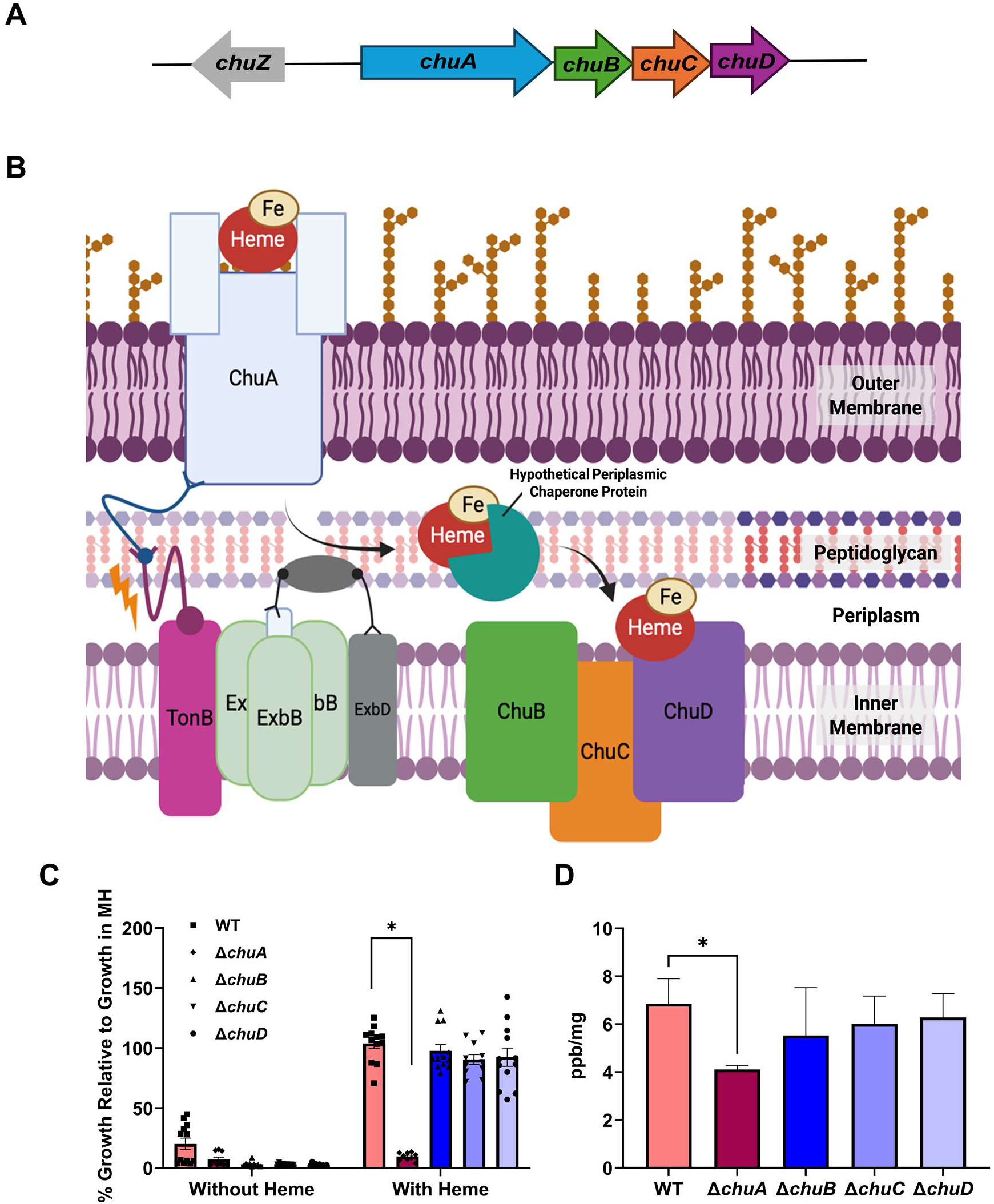
Growth and heme-dependent iron acquisition require ChuA under iron-restriction. (A) Organization of the *chuZABCD* gene cluster. (B) Proposed model of the *C. jejuni* heme transport system, Chu. ChuA is annotated as a TonB-dependent outer membrane heme receptor, ChuB an inner membrane permease, ChuC an ABC transporter, and ChuD a periplasmic hemin-binding protein. In the cytoplasm, the heme oxygenase, ChuZ, processes the heme and releases iron for use in various metabolic processes (not shown). (C) Wild-type, Δ*chuA*, Δ*chuB,* Δ*chuC,* and Δ*chuD* were grown for 48 hours in MH broth with 320 μM DFOM and with 12.5 μM heme or without heme as an iron source. Presented as percent growth when compared to control cultures without DFOM added and are the result of four independent experiments with triplicate cultures. (D) Wild-type, Δ*chuA*, Δ*chuB,* Δ*chuC,* and Δ*chuD* were grown for 48 hours with 320 μM DFOM or without DFOM in the presence of 12.5 μM isotopically labeled heme (^57^Fe-heme) as an iron source, pelleted, washed and weighed. Cell associated ^57^Fe was quantified using ICP-MS and presented. Statistical analysis was performed using unpaired T-tests. *p ≤ 0.05, ****p ≤ 0.0001

Important questions remained following these studies, including whether the Chu system is required for *C. jejuni* infection and what bacterial factors may substitute for the loss of ChuBCD during heme utilization. To address these questions, we created in-frame deletion mutants of ChuA, ChuB, ChuC, and ChuD, and confirmed the earlier results that indicated only ChuA was required for *in vitro* utilization of heme using growth and isotopic iron enrichment assays. Further, we determined that heme utilization gene expression is significantly increased in a murine model of campylobacteriosis and that heme is present in the feces of infected mice and humans. During infection of mice, we observed that wild-type *C. jejuni* and mutants of the ABC transporter efficiently colonized both the cecum and colon of these animals, but that the ChuA mutant was significantly decreased in the colon. This reduction in colonization correlated with significant decreases in neutrophil and monocyte-derived macrophage influx into the colon, which was further associated with a depletion of self-maintaining, embryonically derived tissue-resident macrophages. Further, decreased colonization and innate immune cell recruitment in the colon corresponded with an improvement in multiple markers of tissue pathology, indicating that inhibiting heme acquisition during *C. jejuni* infection may decrease the severity of campylobacteriosis.

## Results

### ChuA is required for heme transport by *C. jejuni* under iron-restricted conditions

To examine which determinants of the heme transport system are required for moving heme into the cell under iron-restricted conditions, we created a series of in-frame deletion mutants: Δ*chuA,* Δ*chuB,* Δ*chuC,* and Δ*chuD.* These strains were grown under iron-restricted conditions either with or without added heme, and their growth was assessed by optical density. When the growth of each strain under iron-restricted conditions without added heme was compared to that under iron-replete conditions (Meuller Hinton (MH) media alone), growth was found to be significantly reduced with mean values of 5.08%, 9.39%, 2.24%, 2.94%, and 2.66% for wild-type, Δ*chuA,* Δ*chuB,* Δ*chuC,* and Δ*chuD,* respectively (Fig. 1C). When heme was added to similar iron-restricted cultures, growth significantly increased for wild-type (107.82%), Δ*chuB* (90.21%), Δ*chuC* (81.97%), and Δ*chuD* (91.63%) relative to iron-replete conditions (Fig. 1C). In contrast, despite the addition of exogenous heme, growth of Δ*chuA* did not significantly increase (10.49%), which could be partially restored by complementation with plasmid-encoded ChuA (Fig. S1). These results suggest that ChuA is required for the transport of heme into *C. jejuni* while the other Chu determinants only modestly affect the use of heme as an alternative iron source under iron-restricted conditions.

To confirm that reduced growth of Δ*chuA* during supplementation was due to decreased transport of heme, we grew similarly restricted cultures of each mutant, added heme containing isotopic iron (^57^Fe), and quantified the relative levels of ^57^Fe in washed and weighed cell pellets by inductively coupled plasma mass spectrometry. From this analysis, we observed that wild-type pellets contained approximately 6.86 ppb ^57^Fe per milligram of biomass. By comparison, Δ*chuA,* Δ*chuB,* Δ*chuC,* and Δ*chuD* cell pellets possessed 4.11, 5.54, 6.01, and 6.29 ppb ^57^Fe/milligram, respectively (Fig. 1D). The 40% decrease in ^57^Fe observed for Δ*chuA* was the only significant reduction among the mutants when compared to wild-type, supporting the growth data that ChuA is the only transport determinant required for heme utilization *in vitro*.

### Heme and transcription of the *C. jejuni* heme transport operon are significantly increased during *C. jejuni* infection

Human campylobacteriosis is known to result in hematochezia, or bloody stool. To confirm heme is present and may serve as an alternative iron source during *C. jejuni* infection, we performed mass spectrometry analysis on the feces of infected humans and mice. To quantify the effect of infection on fecal heme accumulation in humans, we leveraged surveillance stools collected by our group from healthy, uninfected volunteers, as well as clinical samples from patients known to be infected with *Campylobacter,* but no other gastrointestinal pathogen. Using high-performance liquid chromatography (HPLC) to measure fecal heme, we found that feces from uninfected volunteers contained 2.90 pmol/mg and those from infected patients possessed significantly more heme at 753.83 pmol/mg - a 260-fold increase (Fig. 2A). Similarly, analysis of fecal pellets from uninfected or wild-type-infected mice found that heme was significantly elevated by 3.8-fold in the feces of infected mice when compared to uninfected animals (Fig. 2B). These data support the hypothesis that increased fecal heme is consistent with gastrointestinal bleeding during infection, which could promote efficient colonization of the GI tract by *C. jejuni* using the heme transport system.

**Figure 2.**
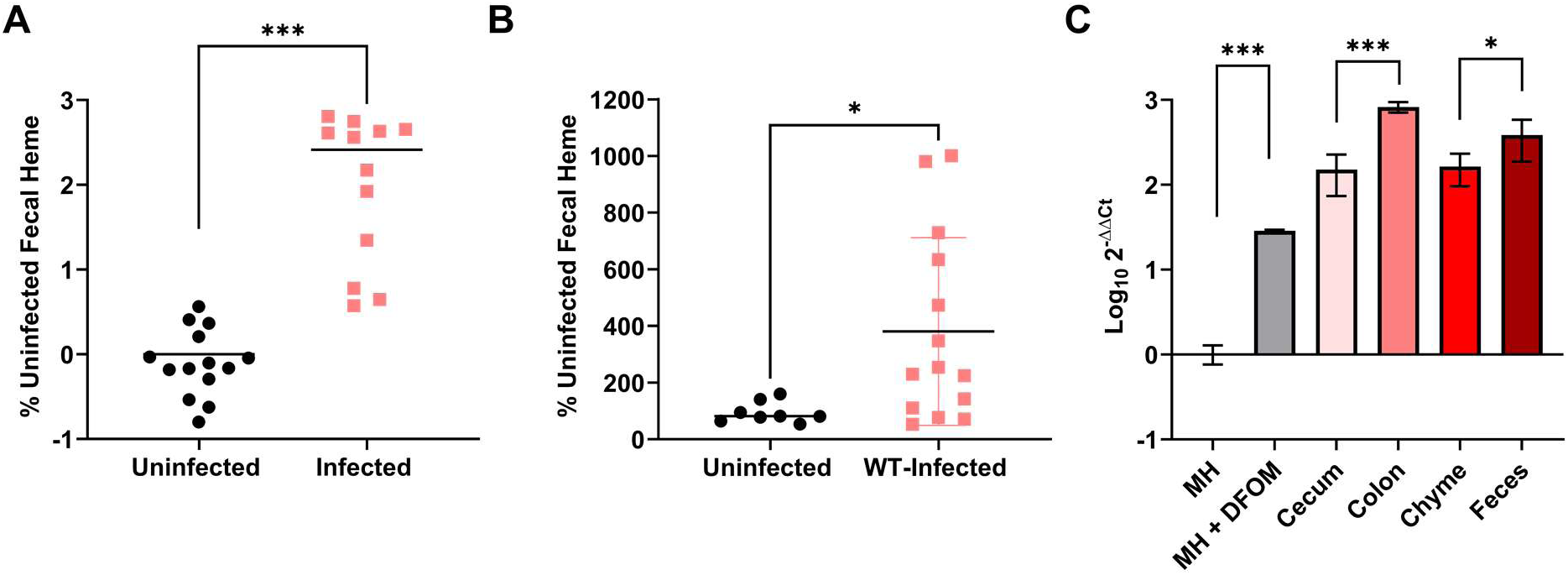
Fecal heme and expression of the Chu heme transport system are abundant during infection. (A) Levels of heme were measured by mass spectrometry in the feces of uninfected volunteers or in clinical samples from patients infected with only *Campylobacter*. (B) Levels of heme b were measured by mass spectrometry in the feces of IL-10^−/−^ mice uninfected (UI) or infected with wild-type *C. jejuni*. (C) RT-qPCR analysis of *chuA* expression in wild-type *C. jejuni* grown in iron-replete MH (MH) or iron-restricted MH (MH+ 320 μM DFOM), or in the cecum, colon, chyme, or feces, during infection of IL-10^−/−^ mice. Expression relative to the *rpoA* housekeeping control. Statistical analysis was performed using Mann-Whitney Test and unpaired T-test. *p ≤ 0.05; **p ≤ 0.01; ***p ≤ 0.001

To examine whether the bacterium expresses the Chu system to acquire heme during murine infection, we extracted RNA from *in vitro* grown iron-replete and iron-restricted media, cecal tissue, colon tissue, cecal contents (chyme), and feces, and quantified *chuA* transcript abundance relative to iron-replete *in vitro* grown cells. From this analysis, we found that *chuA* transcript abundance significantly increased during iron-restriction *in vitro,* with a mean increase of 28.71-fold (Fig. 2C). Similarly, c*huA* abundance was significantly increased in all mouse tissues with fold-changes of 150.48 for cecum, 823.92 for colon, 163.63 for chyme, and 384.77 for feces when compared to iron-replete, *in vitro* grown cells (Fig. 2C). When comparing these values, statistical significance was detected in the 5.48-fold increase of *chuA* in the colon compared to the cecum, as well as the 2.35-fold increase in the feces when compared to the chyme. These changes suggest that during infection, *C. jejuni* increases expression of the Chu system, specifically *chuA,* to gain access to the heme that is released from the host, particularly in the colon and feces.

### ChuA is required for efficient colonization of the colon by *C. jejuni*

Using a murine model of campylobacteriosis, we examined the ability of each heme transport mutant to accumulate within the feces and colonize the cecum and/or colon. All mice were efficiently colonized after a single dose of either wild-type *C. jejuni* or each heme transport mutant with high numbers of the bacterium excreted in the feces 1×10^7^ to 3×10^10^ CFU/g (Fig. 3A). These bacterial loads did not significantly differ between mice infected with either wild-type or the heme transport mutants. After harvesting and plating tissue homogenates for each animal, only Δ*chuC* was found to have significantly reduced colonization in the cecum with a mean tissue load of 1.77×10^7^ CFU/g when compared to 5.83×10^7^ CFU/g for wild-type – a 3.3-fold decrease (Fig. 3B). Similarly, only Δ*chuA,* with a mean load of 5.06×10^6^ CFU/g in the colon, was significantly decreased when compared to wild-type at 3.99×10^7^ CFU/g – a reduction of 7.9-fold (Fig. 3C). Thus, ChuA is dispensable for growth in the intestinal lumen and fecal shedding but is required for efficient colonization of colonic tissue.

**Figure 3.**
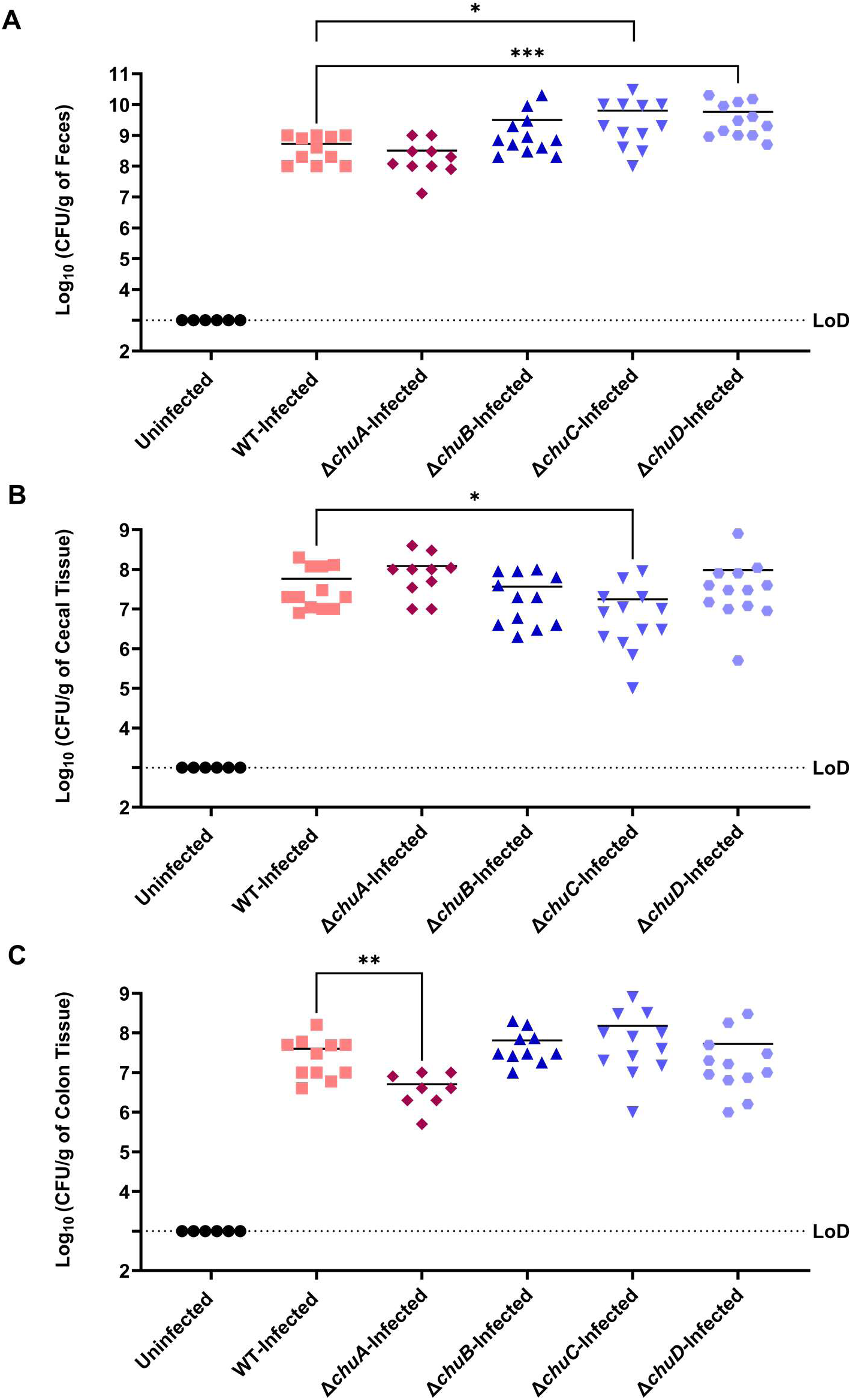
ChuA is required for efficient colonization of the colon. Following *C. jejuni* infection, mice were euthanized, and gastrointestinal tissues were harvested. Colon and cecal contents were removed; the tissue washed and homogenized in PBS. Homogenates were serially diluted, and CFU enumerated following 72 hours of incubation on *Campylobacter*-selective media. Bacterial loads per gram of sample were calculated for feces (A), cecum (B), and colon (C). Statistical analysis was done using a Mann-Whitney test. * p ≤ 0.05, ** p ≤ 0.01, *** p ≤ 0.001, **** p ≤ 0.0001

### Innate immune cell populations in the colon are affected during infection with a ChuA mutant

We quantified neutrophils and macrophages within the ceca and colons of uninfected, wild-type-infected, or *chu* mutant-infected mice after six days of infection using myeloid markers CD11b, Ly6G, and F4/80. To identify neutrophil recruitment, CD11b and Ly6G were used [24]. When examining neutrophils (CD11b/Ly6G double positive cells) in the cecum, there was a significant increase observed for the recruitment of these cells for wild-type-and Δ*chuA*-infected mice when compared to uninfected animals with fold changes of 2.48-and 2.18, respectively (Fig. 4A). Neutrophils within the ceca of Δ*chuB-,* Δ*chuC-,* and Δ*chuD*-infected mice were also increased (Fig. S2A). Infection with wild-type resulted in a significant increase in neutrophils in the colon with an average fold change of 2.01 when compared to uninfected mice (Fig. 4B). Notably, there was not a significant increase of neutrophils in the colons of Δ*chuA*-infected mice with an average fold change of 0.94 when compared to uninfected mice while those infected with Δ*chuB,* Δ*chuC,* or Δ*chuD* remained elevated (Fig. S2B). To quantify mature monocytes and macrophages that were recruited to the site of infection, the myeloid marker CD11b and the monocyte/macrophage marker F4/80 were used [24–26]. When examining the cecum, there was an increase in mature monocytes and monocyte-derived macrophages (CD11b and F4/80 double positive cells) for all infected mice when compared to uninfected animals (Fig. 4C and S2C). Upon infection, there was a significant increase in mature monocytes and macrophages in the colons of wild-type-infected mice when compared to uninfected mice with a fold change of 1.53 (Fig. 4D). Like above, there was not a significant increase observed in the colon for mature monocytes and macrophages in Δ*chuA*-infected mice with a fold change of 0.97 when compared to uninfected mice (Fig. 4D) while those infected with Δ*chuB,* Δ*chuC,* or Δ*chuD* remained higher (Fig. S2D). Lastly, to analyze the number of embryonically derived, self-maintaining tissue macrophages in each organ, the amount of CD11b negative and F4/80 positive cells were quantified [27, 28]. This is a specific subset of macrophages that makes up approximately 20% of the macrophages found in an adult mouse’s gastrointestinal mucosa that are seeded in the GI tract before birth rather than being recruited and differentiated [27, 28]. Wild-type-infected mice exhibited a significant decrease in CD11b^−^/F4/80^+^ tissue-resident macrophages in the cecum and colon when compared to uninfected mice with fold changes of 0.01 and 0.15, respectively (Fig. 4E-F). In contrast, there was not a significant difference in the self-maintaining, embryonically derived tissue macrophage populations between Δ*chuA*-infected and uninfected mice for the cecum and colon with fold changes of 0.63 and 0.42, respectively. Like above, mice infected with Δ*chuB,* Δ*chuC,* and Δ*chuD* had a significant decrease in mature tissue-resident macrophages for the cecum and colon (Fig. S2E-F). These data are consistent with previous findings that infection within the gut, along with other infections, causes an influx of circulating myeloid cells to the tissue and a decrease in tissue-resident cells [29].

**Figure 4.**
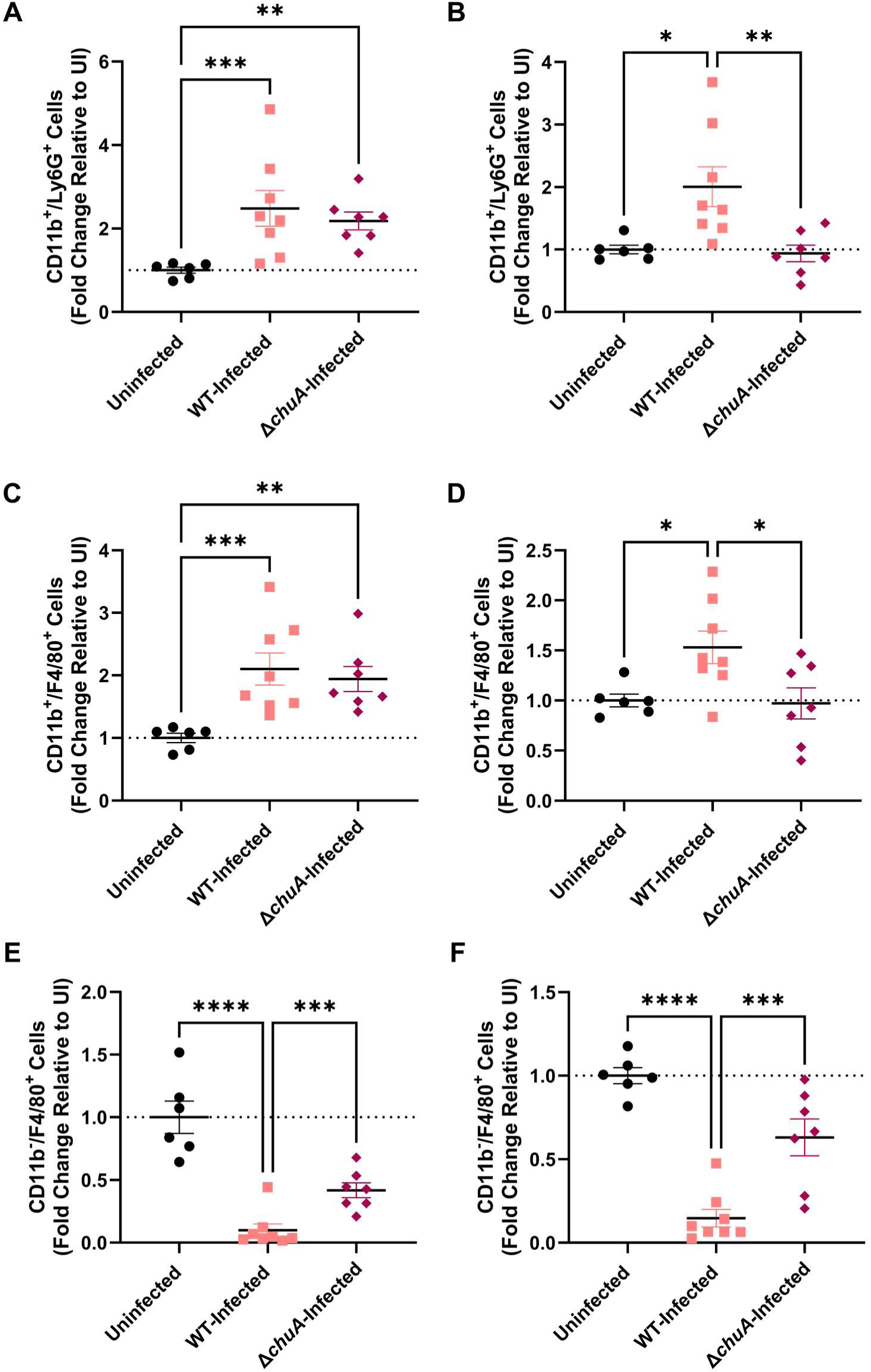
Innate immune cell influx to the colon is dependent on the presence of ChuA. Murine cecal (A, C, & E) and colonic (B, D, & F) tissues from uninfected, wild-type-(WT) infected, and Δ*chuA*-infected mice were harvested post-infection and were homogenized into single cell suspensions for flow cytometry analysis. Cells were incubated with conjugated antibodies for the immune cell surface markers CD11b, Ly6G, and F4/80. Flow cytometry analysis was performed using FlowJo to identify cell populations positive/negative for CD11b and either Ly6G or F4/80 to differentiate between cell types. The percentage of double positive cells was calculated for each population and fold change relative to the same populations in uninfected mice was reported. Statistical analysis was done using a one-way ANOVA with a Tukey’s multiple comparison test. *p ≤ 0.05, **p ≤ 0.01, ***p ≤ 0.001, ****p ≤ 0.0001

### Colon pathology is reduced during infection with a ChuA mutant

To determine whether altered colonization and innate immune cell recruitment affected intestinal tissue pathology, the ceca and colons of infected and uninfected mice were ordinally scored in a blinded manner for apoptotic cells (to ensure any goblet cell loss observed was not due to any cytotoxicity induced by the bacterium), edema, goblet cell loss, and hyperplasia. Representative images for uninfected, wild-type infected, and Δ*chuA* infected colons and ceca are provided (Fig. S5). From this analysis, we determined that the ChuA mutant caused significantly lower goblet cell loss in the cecum when compared to wild-type-infected mice with a score fold change of 1.75 (Fig. 5A). These differences in pathology were not observed during infection with either Δ*chuB,* Δ*chuC*, or Δ*chuD* and there were no differences in apoptotic cells, edema, or hyperplasia noted during infection with any mutant, including Δ*chuA* (Fig. S3AB, Fig. 5B). Further, we also observed lower goblet cell loss and hyperplasia in the colon of Δ*chuA*-infected mice when comparing wild-type-infected mice, with fold changes of 2.90 and 1.75, respectively (Fig. 5C and D). Like above, pathological effects did not significantly differ during infection with Δ*chuB,* Δ*chuC*, or Δ*chuD* and there were no differences observed for apoptotic cells or edema during infection with any mutant, including Δ*chuA* (Fig. S3 C-D). Together, these findings link the ChuA-dependent colonization defect to reduced innate immune cell accumulation and attenuated intestinal pathology.

**Figure 5.**
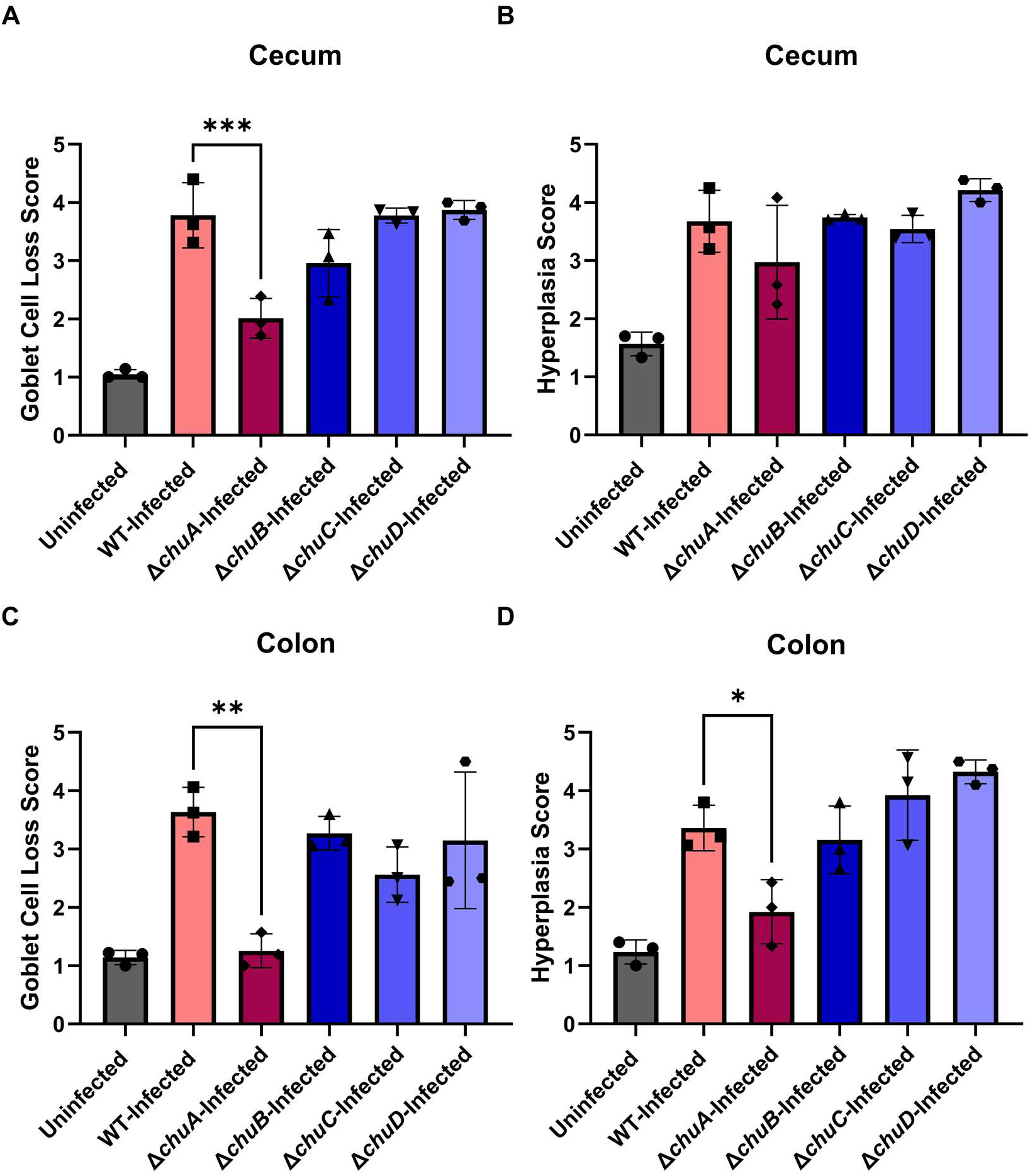
Colon pathology is higher during infection with strains that encode *chuA*. Following infection with wild-type *C. jejuni* or mutant strains, gastrointestinal tissues were formalin-fixed and embedded in paraffin prior to hematoxylin and eosin (H&E) staining. Ceca and colons from three mice in each experimental cohort were examined and blind-scored for goblet cell loss and hyperplasia on a scale of 1-5. Samples were unblinded, and mean scores calculated for cecal goblet cell loss (A), cecal hyperplasia (B), colon goblet cell loss (C), and colon hyperplasia (D). Statistical analysis was done using a one-way ANOVA with a Tukey’s multiple comparison test. *p ≤ 0.05

## Discussion

The host restricts the availability of nutrient metals during infection by limiting access to cofactors required for essential bacterial processes, including respiration and DNA synthesis [13]. Iron represents one of the best-characterized examples of this defense, as it is largely sequestered by host proteins such as hemoglobin, transferrin, and lactoferrin [30–32]. Within the gastrointestinal tract, iron availability is further constrained by competition with the resident microbiota and by complexing with dietary components that are poorly accessible to microbes. These host-imposed limitations have driven the evolution of diverse bacterial iron acquisition systems, including mechanisms to capture, transport, and degrade host-derived heme.

Previous studies examining the transcriptional response of *C. jejuni* during human GI infection reported significant induction of genes encoding the Chu heme uptake system [17]. Consistent with its role in iron acquisition, Chu expression is negatively regulated by Fur and is therefore induced under iron-limiting conditions [18, 33]. Together, these observations suggest that *C. jejuni* experiences iron restriction during infection. Because hematochezia is a common manifestation of campylobacteriosis and represents a potential source of GI heme, we hypothesized that *C. jejuni* uses the Chu system to access host-derived heme and overcome nutritional immunity during intestinal colonization [8].

Consistent with this hypothesis, we found that heme concentrations increased in the GI tracts of infected mice and that *chuA* expression was elevated in both luminal contents and intestinal tissues. Because the gene cluster encoding ChuABCD was previously shown to be operonic, these results suggest the components of heme transport are available in the presence of elevated heme. Surprisingly, only the outer membrane receptor ChuA was required for heme-dependent growth under iron-restricted conditions, whereas mutants lacking the predicted periplasmic and inner membrane transport components retained the ability to utilize heme. This phenotype was recapitulated *in vivo*, where only the *chuA* mutant exhibited a colonization defect, specifically within the colon. Together, these findings indicate that ChuA is the only characterized component of the Chu system required for heme utilization in *C. jejuni* and suggest that transport of heme or heme-derived iron across the inner membrane occurs through an alternative mechanism.

One potential explanation was functional redundancy between ChuBCD and the enterochelin transporter CeuBDE. CeuB and CeuD share modest amino acid similarity with ChuB and ChuC, respectively, and numerous additional ABC transporter ATPases encoded within the *C. jejuni* genome exhibit detectable homology to ChuC. Similar functional overlap between siderophore and heme transport systems has been described in other Gram-negative bacteria, where heterologous ABC transporters can facilitate heme uptake in the absence of their cognate partners [14, 16, 20, 34]. Despite these precedents, disruption of the Ceu transporter failed to compensate for loss of ChuBCD (Fig. S4). These findings leave two intriguing possibilities: either another unidentified ABC transporter mediates inner membrane transport of heme or its degradation products, or *C. jejuni* extracts iron from heme within the periplasm, thereby bypassing the need to transport intact heme across the inner membrane. Distinguishing between these models will be an important direction for future investigation.

Finally, the *chuA* mutant elicited significantly less immune cell influx and pathology in the colon when compared to wild-type *C. jejuni*, which is consistent with its reduced bacterial burden within the tissue. Specifically, infection with the wild-type strain and the *chuBCD* mutants induced robust recruitment of neutrophils, monocytes, and macrophages to the colon, whereas this recruitment was markedly reduced during infection with the *chuA* mutant. Although this decrease in pathology is likely a consequence of impaired colonization, we cannot exclude the possibility that the loss of the surface-exposed ChuA receptor also alters bacterial interactions with the host immune system. The above inflammatory response was also accompanied by a reduction in the self-maintaining tissue-resident macrophages that are of embryonic origin rather than the recruited, bone-marrow derived macrophages, a feature commonly associated with intestinal inflammation [29]. This finding is consistent with a model where resident and self-maintaining macrophages are depleted during infection while recruited monocytes remain in an inflammatory state rather than differentiating into new tissue-resident macrophages.

An additional finding was that the colonization defect of the *chuA* mutant was restricted to the colon, despite induction of *chuA* expression in both the cecum and colon. Notably, *chuA* expression was substantially higher in the colon and feces. This suggests that iron limitation is more severe in the distal intestine, and that dependence on heme acquisition increases accordingly. While it remains unclear why heme acquisition may be needed in the colon, this regional difference may reflect increased microbial competition for iron, progressive depletion of accessible iron along the intestinal tract, or greater iron demands by *C. jejuni* during colonic colonization. Overall, these findings identify ChuA-dependent heme acquisition as a niche-specific adaptation that promotes efficient colonization of the colon and the associated inflammatory pathology during *C. jejuni* infection.

## Material and Methods

### Bacterial Strains and Growth Conditions

All *C. jejuni* strains, including *C. jejuni* 81-176 DRH212 (wild-type) strain and its mutants that were used in this study, were stored at −80°C in Mueller-Hinton (MH) broth supplemented with 20% glycerol. *C. jejuni* strains were routinely cultured on MH agar supplemented with 10% sheep blood and 10 μg/mL trimethoprim (TMP) under microaerobic conditions (85% N_2_, 10% CO_2_, 5% O_2_) for 24-72 hours. *Escherichia coli* strains were stored at −80°C in low-salt LB broth supplemented with 20% glycerol and were routinely grown aerobically at 37°C in LB broth or on LB agar plates, with ampicillin (100 μg/mL) alone or with both ampicillin and chloramphenicol (30 μg/mL) added when required. A complete list of strains used in this study is provided in Supplementary Table S1.

### Mutant Generation

Mutants of *C. jejuni* 81-176 DRH212 were constructed using a two-step allelic exchange approach that began with splicing overlap extension PCR (SOE-PCR). For each target gene of *chuABCD* and *ceuBDE* operon, approximately 500 bp of the upstream and downstream flanking regions of each gene were amplified by PCR and stitched together via SOE-PCR. The resulting fragment was cloned into the pJET1.2/blunt vector according to the manufacturer’s protocol (Thermo-Fisher # K1231). Transformants were selected on LB agar supplemented with ampicillin (100 μg/mL) and screened by colony PCR, followed by sequencing. Confirmed plasmids were digested with BamHI, and a BamHI-digested *rpsL-cat* cassette was ligated between the flanking regions. The resulting constructs were verified by colony PCR and sequencing. These plasmids were electroporated into wild-type *C. jejuni* and chloramphenicol-resistant (Cmᴿ, 30 μg/mL) transformants were selected. Intermediate insertion-deletion mutants (e.g., Δ*chuA*::*rpsL-cat,* Δ*ceuB*::*rpsL-cat*) were first obtained. Clean, in-frame deletion mutants were then generated by electroporating the original SOE-PCR fragment (lacking the *rpsL*::*cat* cassette) into the intermediate strains, followed by selection for chloramphenicol-sensitive and streptomycin-resistant colonies. Correct in-frame deletions were confirmed by PCR and whole-genome sequencing.

Double mutants were constructed by electroporating the *ceuB/D/E*::*rpsL-cat* plasmid into competent cells of the clean Δ*chuB*, Δ*chuC*, or Δ*chuD* single-deletion mutants. Chloramphenicol-resistant transformants were selected and verified by PCR and whole-genome sequencing to confirm the presence of both mutations, resulting in the double mutants Δ*chuB*Δ*ceuB::rpsL-cat*, Δ*chuC*Δ*ceuD*::*rpsL-cat*, and Δ*chuD* Δ*ceuE*:*:rpsL-cat*. Several attempts were made to generate clean, in-frame deletion of double mutants. This was successful only for the Δ*chuD*Δ*ceuE* double mutant; attempts to obtain clean in-frame deletions of the Δ*chuB*Δ*ceuB*::*rpsL-cat* and Δ*chuC* Δ*ceuD*::*rpsL-cat* double mutants were unsuccessful. A complete list of all strains and mutants generated in this study is provided in Supplementary Table S1.

### Δ*chuA+* pRY112*chuA* Construction

Wild-type *C. jejuni* 81-176 was grown on MH + TMP (10 μg/mL) agar plates under microaerobic conditions and genomic DNA was extracted. The *chuA* gene, including its native promoter, was amplified by PCR using primers listed in Supplementary Table S2 and gel extracted. The purified *chuA* fragment was then assembled into the shuttle vector pRY112 using Gibson Assembly according to the manufacturer’s instructions (Gibson Assembly Master Mix) (NEB #E2611). The assembled product was transformed into chemically competent *E. coli* DH5α cells (Thermo Scientific #18258012), and transformants were selected on LB agar plates containing 30 μg/mL chloramphenicol. Resistant colonies likely containing the pRY112*chu*A construct were screened by PCR and confirmed by sequencing. pRY112*chu*A was extracted and electroporated into *E. coli* DH5α pRK212.1 cells. Conjugation was performed between the *E. coli* DH5α pRK212.1 carrying pRY112*chu*A (donor strain) and the Δ*chuA* mutant (recipient strain). Briefly, overnight cultures of the donor and recipient were mixed and spotted onto non-selective MH agar plates, then incubated under microaerobic conditions at 37°C for 5 hours. After incubation, bacteria were resuspended in MH broth, pelleted, and plated onto MH agar containing TMP (10 μg/mL) and chloramphenicol (30 μg/mL). Plates were incubated for 3–5 days under microaerophilic conditions at 37°C. Eight transconjugants were picked after 5 days of incubation and streaked for purity. Genomic DNA was extracted from the transconjugant colonies, and successful transfer and presence of the *chuA* gene were confirmed by PCR.

### Growth Assay

Wild-type *C. jejuni* 81-176 Sm^R^ (DRH212), all the mutant strains and Δ*chuA+* pRY112*chuA* were grown on MH +TMP (10µg/mL) agar plates by spot inoculation for 24 hours, followed by quadrant streaking onto fresh MH agar plates for an additional 24 hours under microaerobic conditions at 37°C. Bacterial cells were resuspended in MH broth to an optical density at 600nm (OD A600) of 0.05. 100 μL of this suspension was added to wells containing 100μL of MH broth supplemented with either 640 μM deferoxamine (DFOM) alone, 640μM DFOM plus 25μM heme or MH only. This resulted in a final volume of 200μL per well with final concentrations of 320 μM DFOM, 320 μM DFOM plus 12.5 μM heme, or MH only, and a starting OD A600 of 0.025. Plates were incubated microaerobically at 37°C, and OD A600 was measured at 0, 24, and 48 hours. Data from the 48-hour time point were analyzed and presented. Outliers were removed using the robust regression and outlier removal (ROUT) method with Q=1%.

### ⁵⁷Fe-Heme Growth Assay

*C. jejuni* strains were grown on MH + TMP media as above and used to inoculate 5 mL overnight cultures in MH broth. These cultures were then used to inoculate 5 mL fresh MH broth in optically clear tubes to a starting OD A600 of 0.025 under four conditions: i) MH alone, ii) MH with 320 µM DFOM, iii) MH with 12.5 μM ^57^Fe-heme (Fisher Specialty Chemicals, Cat. # P40080), or iv) MH with 12.5 μM ^57^Fe-heme and 320 µM DFOM. Cultures were incubated microaerobically at 37°C until mid-logarithmic phase (approximately 30 hours). OD A600 was measured for each condition across all strains. Cells were harvested by centrifugation at 3000 x g for 8 minutes, washed twice with sterile water, resuspended in 500 µL sterile water, and weighed. The bacterial pellets were sent to Vanderbilt University for ICP-MS analysis.

The samples were transferred to a 15mL metal-free tube, where 2x volume of nitric acid was added to digest overnight at 50°C. Then a 3x volume of MilliQ water was added to dilute the nitric acid concentration to a level the instrument could handle. For a total dilution factor of 12x which was accounted for in the data processing. The samples were run on an Agilent 7700 ICP, and data was analyzed with the Mass Hunter Software. The optimal instrument parameters are determined on the day of the run; the samples were initially taken up at 0.5 rps for 30 seconds, followed by 30 seconds at 0.1 rps to stabilize the signal. Spectrum mode analysis was performed at 0.1 revolutions per second, collecting three points across each peak and conducting three replicates of 100 sweeps for each element. The sampling probe and tubing were rinsed with 2% nitric acid for 30 seconds at 0.5 rps between each sample.

### RNA Extraction and Chu Gene Expression

A segment of each tissue, feces, or cecal content (chyme) was placed in 1 mL of cold TRIzol reagent (Invitrogen #15596026) and homogenized. Phase separation and RNA isolation were then performed following the reagent user guide provided by the manufacturer. *C. jejuni* RNA was isolated by spinning down cultures grown in MH +/− DFOM at max speed for 10 minutes at 4°C, removing the supernatant, and resuspending the pellet in 1mL of cold TRIzol reagent. Phase separation and RNA isolation were then performed according to the reagent user guide. Following RNA extraction, cDNA was synthesized using the Bio-Rad iScript cDNA Synthesis Kit (BioRad #1708891). The cDNA library was diluted 1:20 with water, and 5 μL was used in each 20uL qPCR reaction. qPCR was performed using the BioRad iTaq Universal SYBR Green Supermix kit (BioRad #1725121) following the manufacturer manual.

### Heme Quantification in Mice and Human

Fecal pellets were collected on Day 0 (pre-inoculation) and Day 6 post-infection from mice infected with wild-type *C. jejuni* 81-176 and uninfected controls. Pellets were sent to the Louisiana State University School of Veterinary Medicine Mass Spectrometry Resource Center for heme quantification. Fecal pellets were suspended in PBS to a volume of 200 µL. Heme was extracted thrice with 1 mL of 1% acetic acid in ethyl acetate. The fecal pellet was dispersed and exposed to organic solvent through extensive vortexing. Organic/aqueous solvent phase separation was expedited by tabletop centrifugation at 4000 rpm and 4°C for 5 minutes. The ethyl acetate layers were transferred to clean glass tubes, combined, and dried to completion under a stream of nitrogen gas. Each sample was dissolved in 100 µL of methanol for LC-MS/MS analysis. A standard mixture of heme was prepared in methanol at 5, 0.5, 0.05, and 0.005 µM to serve as an external calibration line for quantification.

The samples were analyzed at the LSU School of Veterinary Medicine Mass Spectrometry Resource Center on a Shimadzu 8060NX triple quadrupole mass spectrometer interfaced with a Shimadzu Nexera XS 40 series UHPLC and Shimadzu CTO-40S column oven. 10 µL of each sample, blank, or standard mixture was analyzed in positive ionization mode. Chromatographic separation was achieved using a Shimadzu Nexcol C_18_ (50 mm length, 2.1 mm internal diameter, 1.8 mm particle size) column with a Phenomenex Security Guard C_18_ (2.1 mm internal diameter) guard column. Mobile phase A was 0.1%formic acid in water, and mobile phase B was 0.1% formic acid in acetonitrile. The column oven was set to 25°C, and the flow rate was set to 0.4 mL/minute for the entire analysis. The starting condition was 40% B, which was held for 1 minute. The B mobile phase was then ramped to 98% over the next 14 minutes. The column was washed at 98% B for 5 minutes, then equilibrated to 40% B for 3 minutes. All instrument voltages were determined and optimized empirically prior to the analysis. The m/z transitions were 616.3 to 557.3 for heme. An external calibration line of heme was used to quantify each sample before normalizing total fecal pellet weight. Statistical analysis was performed using a Mann-Whitney U-test.

### Mouse Infection

Eight-to ten-week-old female IL-10^−/−^ C57BL/6 mice were first treated with a broad-spectrum antibiotic mixture consisting of ampicillin (1 g/L), neomycin (10 mg/mL), vancomycin (5 mg/mL), metronidazole (10 mg/mL), and amphotericin B (0.1 mg/mL). The antibiotic mixture was administered at a dose of 10 µL per gram of body weight via oral gavage at 12-hour intervals for 7 days. Following the final antibiotic dose, a 72-hour washout period was observed to allow clearance of residual antibiotics. Subsequently, wild-type *C. jejuni* strain 81-176 and Δ*chu* mutant strains were cultured on *Campylobacter*-selective (CS) medium containing 10% sheep’s blood, cefoperazone (40 µg/ml), cycloheximide (100 µg/ml), trimethoprim (10 µg/ml), and vancomycin (100µg/ml) agar plates and Gram stained to verify culture purity. Fresh bacterial suspensions were prepared in sterile 1× PBS at a concentration of approximately 10^10^ CFU/mL. Mice were then inoculated with a sterile 1x PBS control, 10^9^ CFU of wild-type *C. jejuni* 81-176, or 10^9^ CFU of one of the Δ*chu* mutant strains by oral gavage. Fecal pellets were collected on day 6 post-infection, weighed, and homogenized in sterile 1x PBS at a 1:10 (w/v) dilution. The resulting homogenates were serially diluted and plated onto CS media. Plates were incubated at 37°C under microaerobic conditions for 72 hours, after which CFU were enumerated. On day 6 post-infection, mice were humanely euthanized, and the cecum and colon were harvested and washed in sterile 1× PBS. Approximately one third of each tissue sample was weighed, diluted 1:100 (w/v) in 1x PBS, homogenized, serially diluted, and plated on CS media. Plates were incubated for 72 hours microaerobically at 37°C, and the CFU were counted.

### Histopathology Scoring

Following euthanasia of IL-10^−/−^ C57BL/6 mice, the cecum and colon were harvested. The organs were separated, sliced longitudinally, rinsed with PBS, and then approximately 2/3 of each organ was Swiss rolled. The rolled organs were stored in 10% formalin for 48 hours, then underwent a progressive series of alcohol and xylene dehydrations. The rolled organs were embedded in paraffin, sectioned (∼5 µm), and mounted onto super frost plus microscope slides (Fisher Scientific #1255015). The slides were routinely stained with hematoxylin and eosin (HE) to evaluate histopathology and assign tissue scoring. For the scoring, images were captured at 20X objective in color brightfield mode using an Agilent BioTek Cytation 5 cell imaging reader (Agilent Technologies, Santa Clara, CA, USA). All slides were coded and evaluated in a manner blinded to the group assignment to prevent observer bias [35]. For each mouse, at least 10 non-overlapping microscopic fields were scored for both cecum and colon. Edema, hyperplasia, and goblet cell loss were evaluated in each field using an ordinal scoring system on a scale 1 to 5 where 1 indicates normal appearance and 5 indicates severe pathology. Apoptotic bodies observed in each field were counted for each mouse to assess any potential loss of goblet cells to cytotoxicity. For each parameter, the scores from all scored fields were summed and divided by the number of fields to obtain the mean score per mouse. After completing all the scoring, the samples were unmasked to group assignments. Data was graphed and statistically analyzed using ordinary one-way ANOVA.

### Flow Cytometry

Following euthanasia, mice were necropsied, and the cecum and colon from each mouse were removed and placed in sterile 1x PBS. Fecal pellets were removed from the colon and saved for colonization plating. The organs were then cut longitudinally and washed 1x PBS to clean the tissue of any residual cecal/fecal content. Approximately one third of the tissue was removed and saved for colonization plating and the rest was prepped for flow cytometry analysis. Each tissue section in pre-weighed tubes with 1x PBS was weighed and then poured over a 70 μM cell strainer (Corning #0052) in a 50 mL conical tube. The tissue was grounded into the cell strainer using a 10 mL syringe plunger. The PBS in the conical tube was used to carefully wash the cell strainer at least two times to remove any cells stuck in the strainer. The PBS/organ suspensions were centrifuged at 800 x g for 10 minutes at 4°C, and the supernatant was then removed. Pellets were resuspended in 1 mL of 1x RBC lysis buffer (BioLegend #420301) and incubated at room temperature for 8 minutes. Following incubation, 10 mL of fresh 1x PBS was added. The liquid was removed using a serological pipette and ran through a new cell strainer into a new conical tube. Cell suspensions were centrifuged again at the same settings as the first spin; supernatant was removed, and the pellets were resuspended in 800 μL of FACS buffer (1x PBS with 2% HI FBS and 0.02% sodium azide). 200 μL of each suspension was pipetted into a 96 well U-bottom plate in triplicate and then prepared for flow cytometry as described below.

After aliquoting each suspension to the appropriate wells in 96 well plates, plates were centrifuged at 800 x g for 10 minutes at 4°C. Following centrifugation, supernatant was removed by flicking the plate into a waste container, and the pellets were immediately fixed by resuspending in 100 μL of 1x PBS with a final concentration of 4% paraformaldehyde (Thermo Scientific #28908). This was incubated on ice for 20 minutes, and the plates were centrifuged with the settings used previously. Following centrifugation, supernatant was removed, and pellets were resuspended in 100 μL of FACS buffer with FC block added at a 1:150 concentration (TonboBiosciences Anti-Mouse CD16/CD32 (Fc Shield) #70-0161-M001). Plates were incubated on ice for 20 minutes, centrifuged, and supernatant was removed as previously described. Cell pellets were then resuspended in 800 μL of FACS buffer containing the conjugated primary antibodies, each at a 1:800 concentration (BioLegend APC anti-mouse/human CD11b #101212, BioLegend PerCP anti-mouse/human Ly-6G #127654, BioLegend Brilliant Violet 711 anti-mouse F4/80 #123147). These were incubated on ice for 20 minutes in the dark, centrifuged, and supernatant was removed. Each cell pellet was resuspended in 250 μL of FACS buffer and analyzed using a Cytek Aurora Flow Cytometer.

### Flow Cytometry Analysis

Flow cytometry data were analyzed using the FlowJo Software. After uploading all .fcs files, doublets and cell clumps were gated out using FSC-H vs FSC-A. Single cells were then gated on single or double positive signal for each fluorescent marker based on comparison to single and FMO stained controls. Percentages of each cell type were calculated using the number of positive cells and the total number of single cells. These percentages were normalized to uninfected mice from the same experiment and reported as fold changes relative to uninfected. Statistical analysis was performed using ordinary and lognormal ordinary one-way ANOVA tests.

## Acknowledgements

The research reported in this publication was supported by University of Iowa start-up funds provided to J.G.J., the Carver College of Medicine Stead Family Innovation Scholars Program awarded to J.G.J., NIH funds R01AI166535 to J.G.J., K22AI153677 to W.N.B., R35GM154838 to A.J.M. and R35GM154857 to W.N.B. Research was also supported by USDA funds to J.G.J.: NIFA-AFRI-2019-67017-29261. The University of Tennessee provided funds to A.J.M., including start-up and Human Health and Wellness Initiative funds. The authors would like to acknowledge use of the University of Iowa Central Microscopy Research Facility, a core resource supported by the University of Iowa Vice President for Research, and the Carver College of Medicine. The flow cytometry data presented herein were obtained at the Flow Cytometry Facility, which is a Carver College of Medicine / Holden Comprehensive Cancer Center core research facility at the University of Iowa. The facility is funded through user fees and the generous financial support of the Carver College of Medicine, Holden Comprehensive Cancer Center, and Iowa City Veteran’s Administration Medical Center. Flow cytometry research reported in this publication was supported by the Carver Charitable Trust - Grant #26-6110. The authors would like to thank Dr. Wade Calcutt and Brian Hachey at the Vanderbilt University Mass Spectrometry Research Center for their assistance with the various analyses.

---

**Figure S1.**
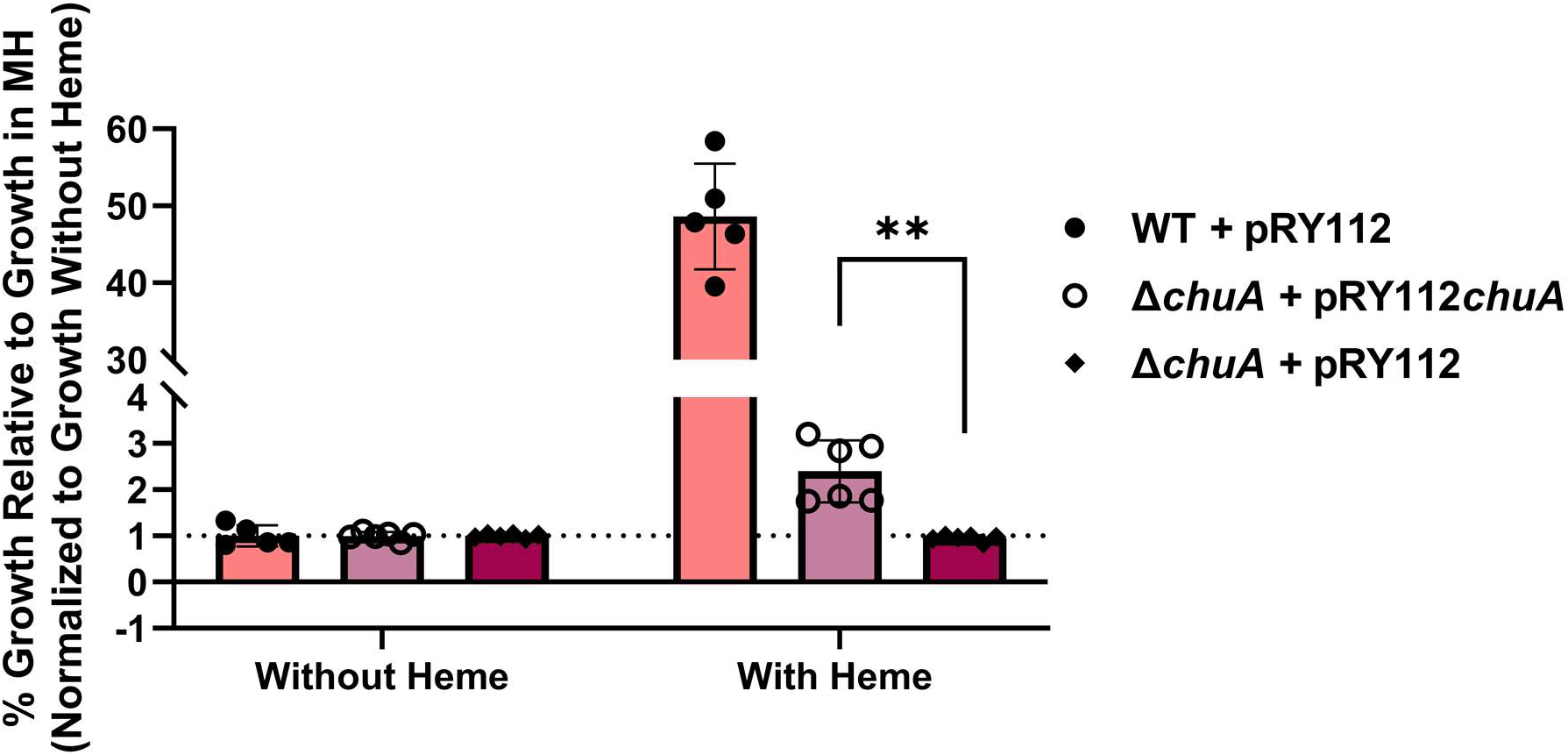
Complementation of the Δ*chuA* mutant with *chuA* restores heme-dependent iron acquisition and growth. Wild-type (WT), Δ*chuA* mutant and Δ*chuA* + pRY112*chuA* strains were grown for 48 hours in MH broth containing 320 μM DFOM and with 12.5 μM heme or without heme supplementation. Data are presented as growth relative to growth in MH + 320 μM DFOM and are the result of two independent experiments with triplicate cultures. Statistical analysis was done using unpaired two-tailed Welch’s t-test. **p ≤ 0.01

**Figure S2.**
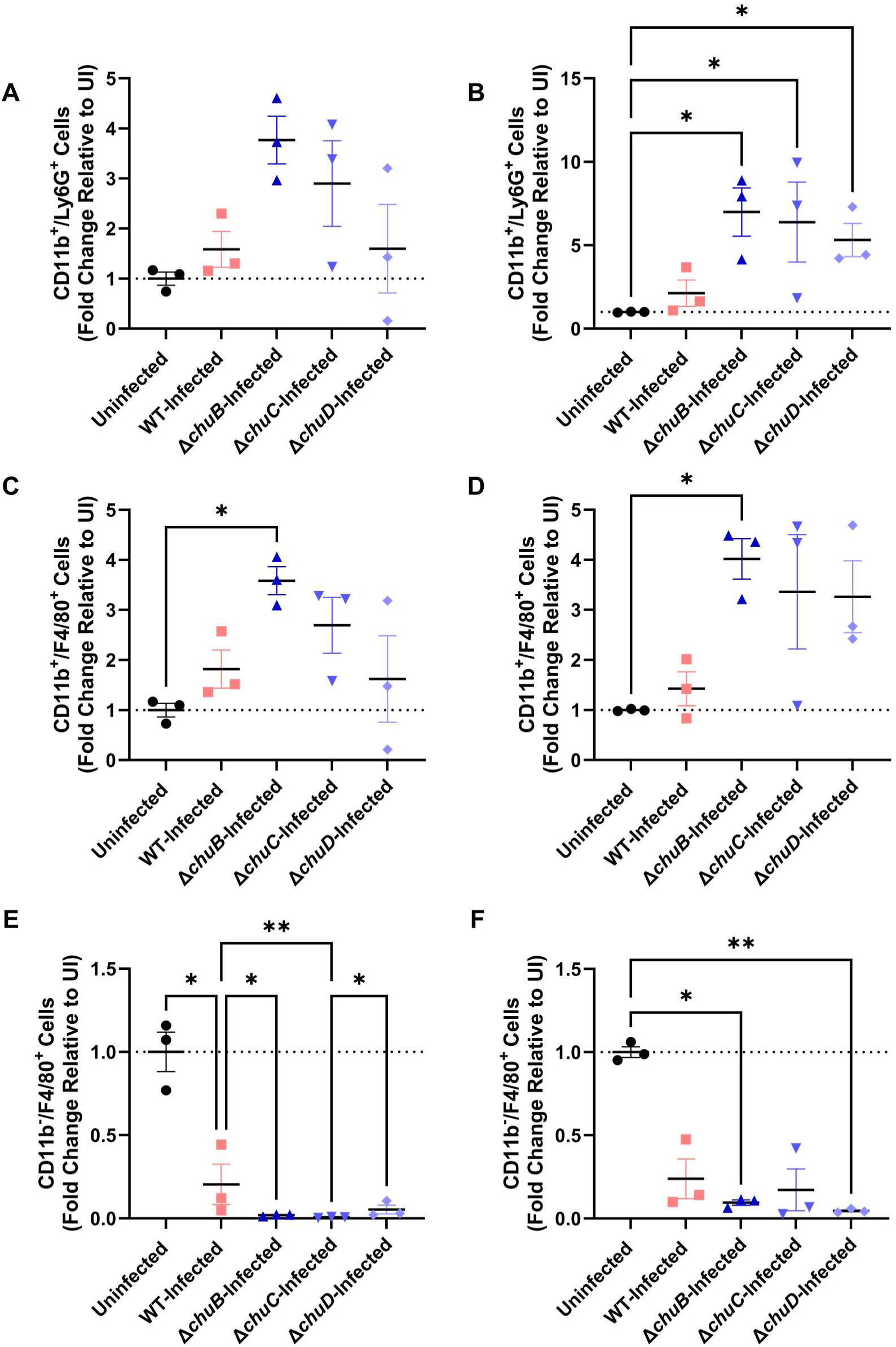
Innate immune cell populations are recruited in levels similar to wild-type when infected with mutants in ChuB, C, or. **D.** Murine cecal (A, C, & E) and colonic (B, D, & F) tissues from uninfected, wild-type-(WT) infected, and Δ*chuB-,* Δ*chuC-,* and Δ*chuD*-infected mice were harvested post-infection and were homogenized into single cell suspensions for flow cytometry analysis. Cells were incubated with conjugated antibodies for the immune cell surface markers CD11b, Ly6G, and F4/80. Flow cytometry analysis was performed using FlowJo to identify cell populations positive/negative for CD11b and either Ly6G or F4/80 to differentiate between cell types. The percentage of double positive cells was calculated for each population and fold change relative to the same populations in uninfected mice was reported. Statistical analysis was done using a one-way ANOVA with a Tukey’s multiple comparison test. *p ≤ 0.05, **p ≤ 0.01, ***p ≤ 0.001, ****p ≤ 0.0001

**Figure S3.**
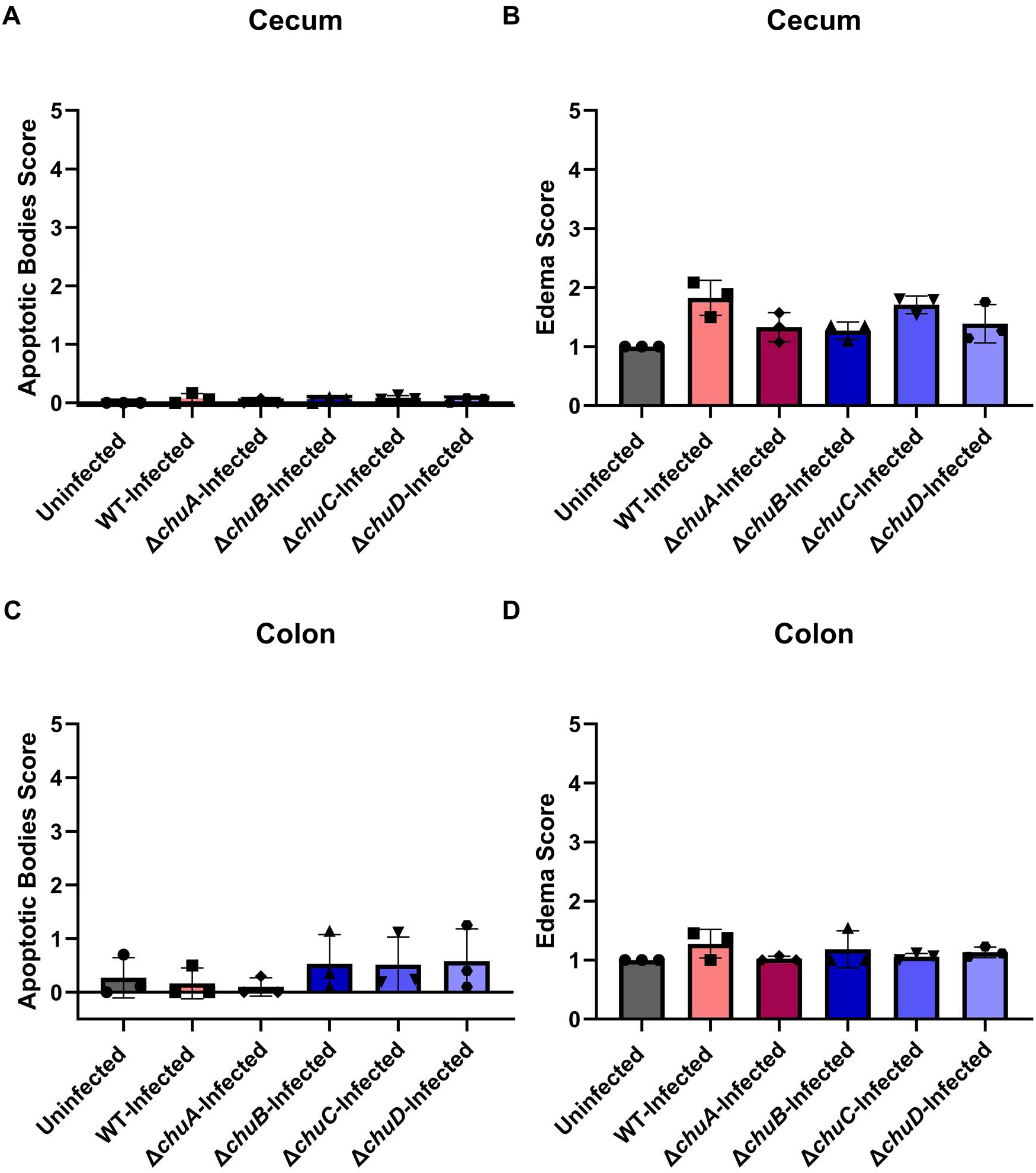
Infection with the Δ*chuA* mutant does not induce significant apoptosis or edema in the cecum and colon. Mice were infected with the *C. jejuni* wild-type (WT) strain or one of the mutant strains. Ceca and colons from three mice in each experimental cohort were examined and blind-scored for cecal apoptotic body scores (A), cecal edema scores (B), colon apoptotic body scores (C), and colon edema scores (D). Apoptotic body score was determined by counting apoptotic bodies across all fields and averaging the total per field. Edema was scored across all fields on a scale of 1 to 5, where 1 represents normal tissue, and 5 represents severely inflamed tissue, and then averaged per field. All tissue sections were evaluated under a 20x objective lens. Statistical analysis was done using a one-way ANOVA with a Tukey’s multiple comparison test. * p ≤ 0.05, ** p ≤ 0.01

**Figure S4.**
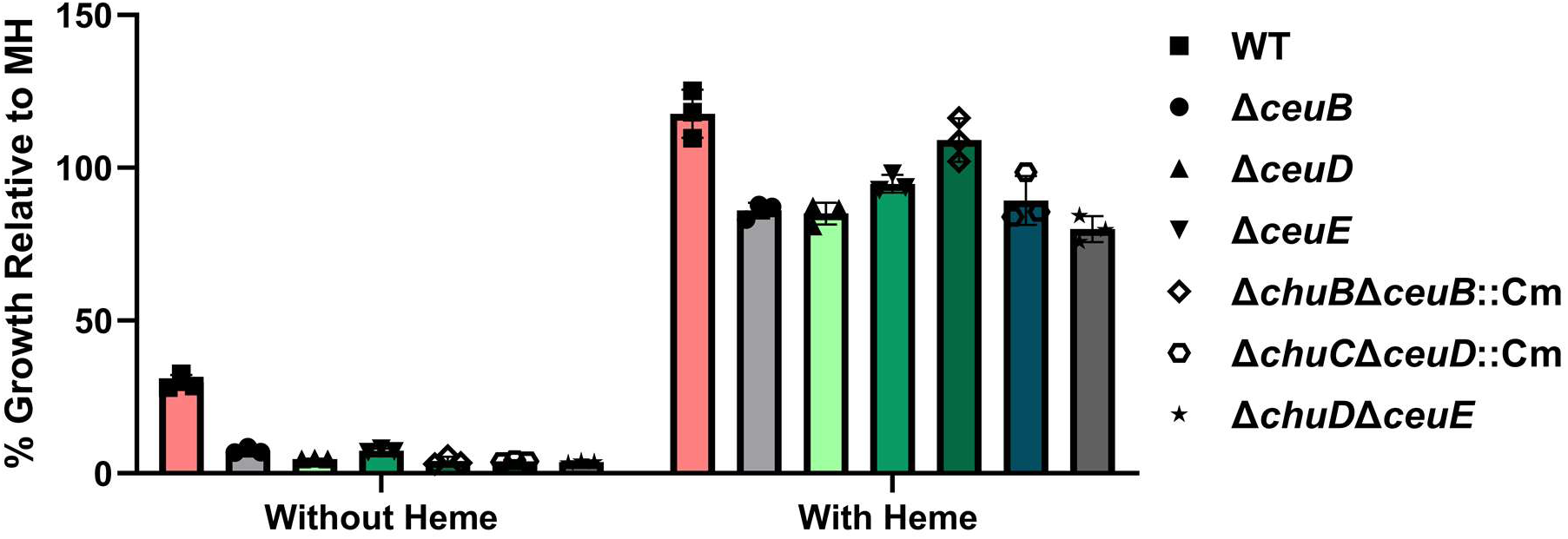
*ceuBDE* and *chuBCD*-*ceuBDE* systems are dispensable for heme-dependent growth of *C. jejuni* under iron-restricted conditions. Wild-type (WT), Δ*ceuB,* Δ*ceuD,* Δ*ceuE,* Δ*chuB*Δ*ceuB*::Cm, Δ*chuC*Δ*ceuD:*:Cm and, Δ*chuD*Δ*ceuE* mutant strains were grown for 48 hours in MH broth containing 320μM DFOM and with 12.5μM heme or without heme supplementation. Growth was measured as optical density and expressed as percentage relative to growth in MH. Data presented is representative of two independent experiments, with triplicate cultures per experiment.

**Figure S5.**
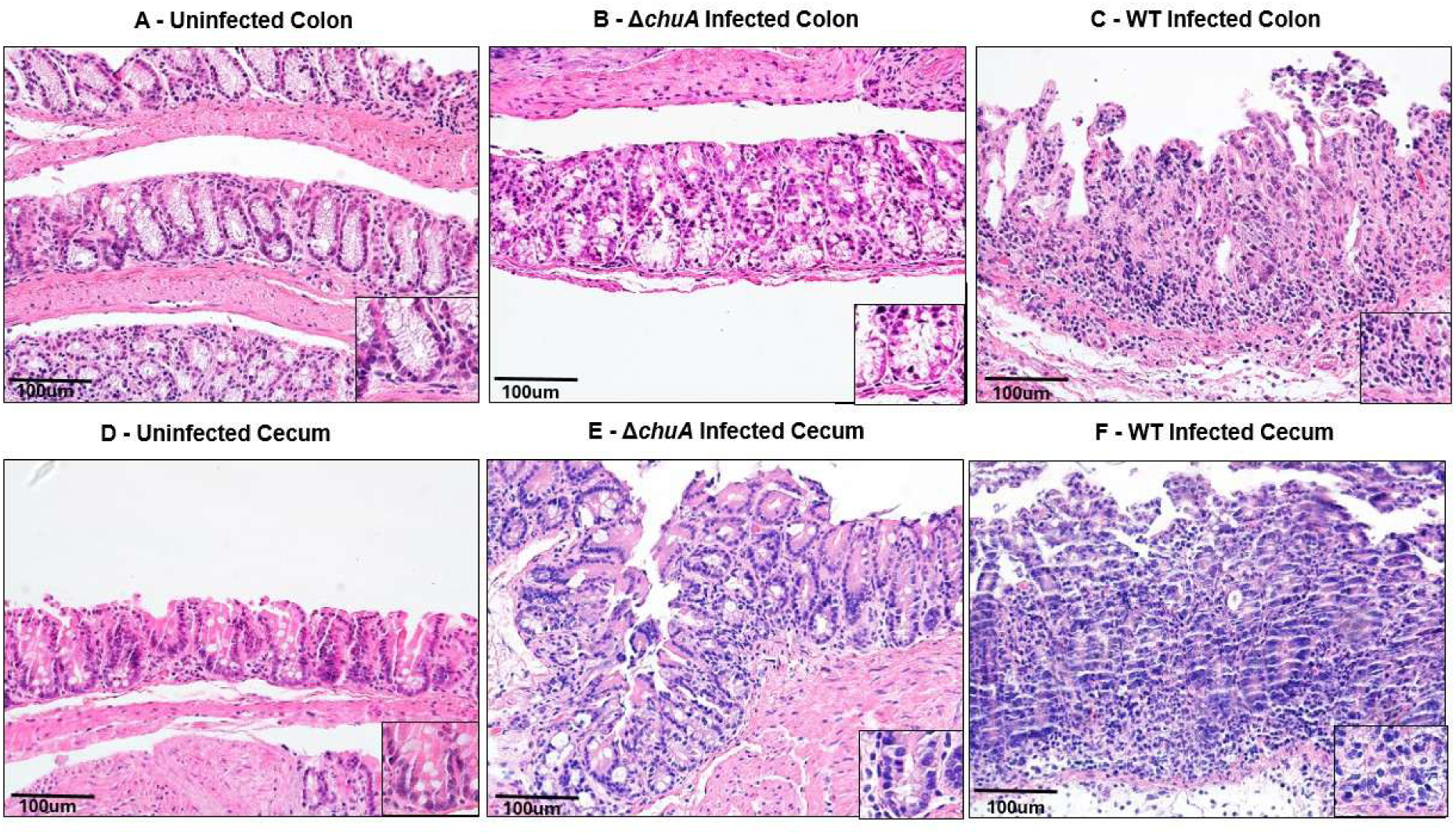
Representative H&E-stained sections of the colon and cecum. Representative hematoxylin and eosin-(HE) stained sections of the colon from (A) uninfected, (B) Δ*chuA* mutant-infected, and (C) wild-type-(WT) infected mice, and of the cecum from (D) uninfected, (E) Δ*chuA*-infected, and (F) WT-infected mice. Images were acquired using a 20X objective. Scale bar, 100 µm.

**Table S1.**
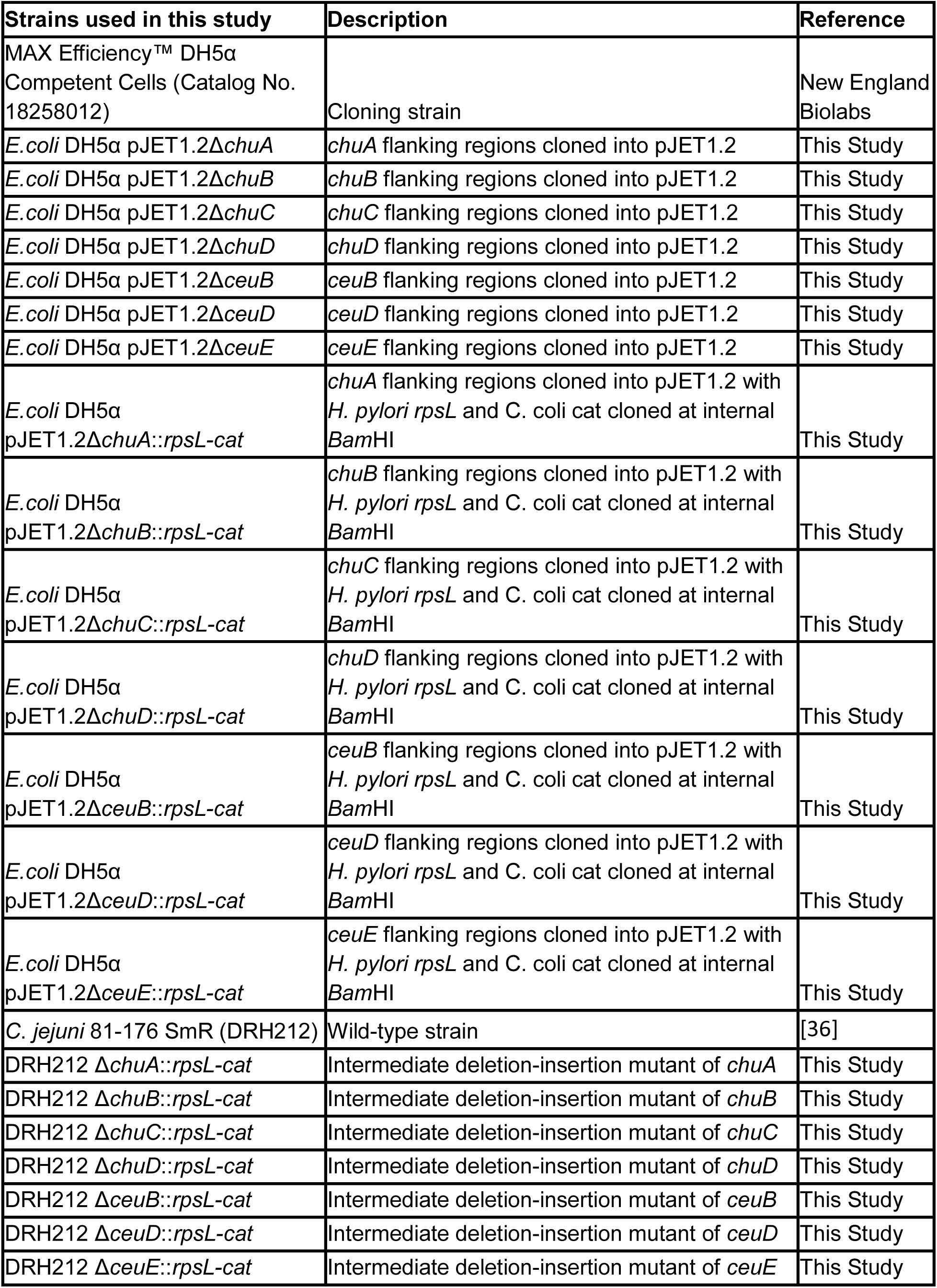
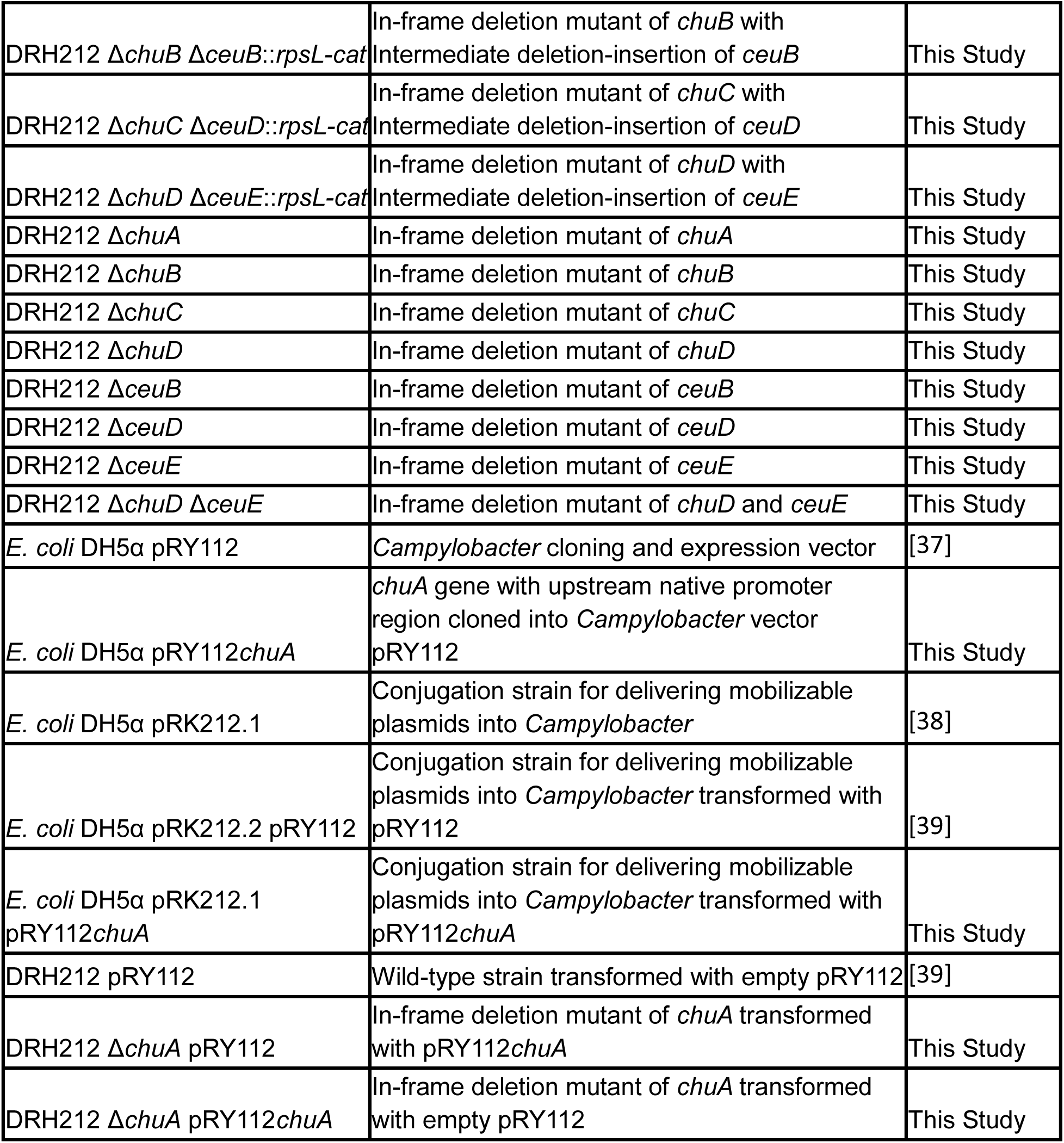
Strains used in this study.

**Table S2.**
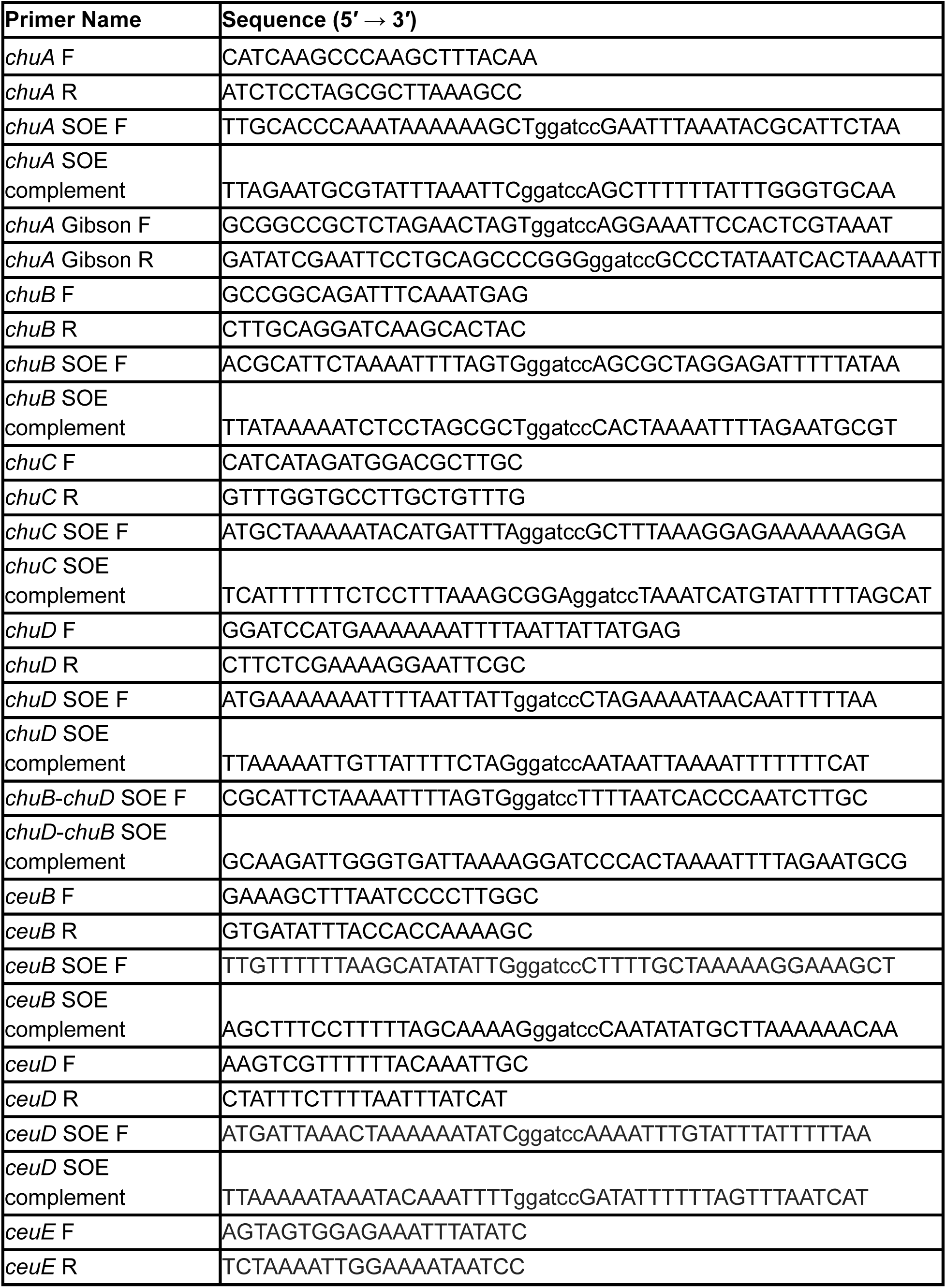
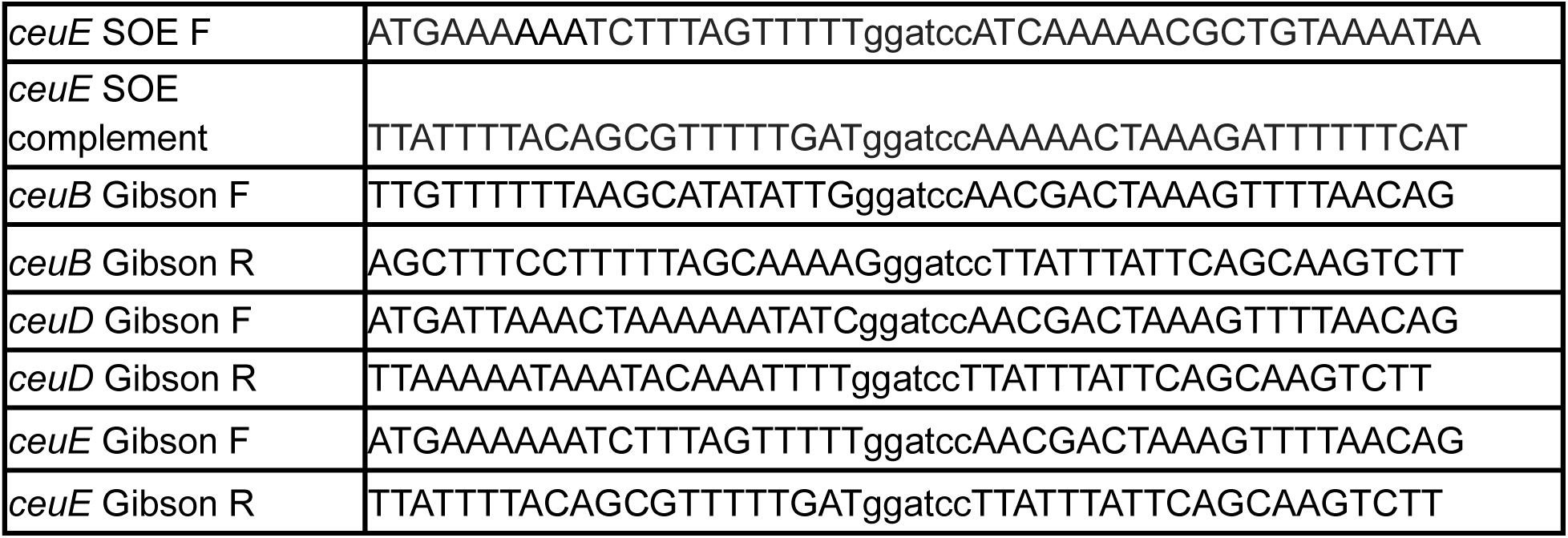
Primers used in this study.

